# Puzzle Hi-C: an accurate scaffolding software

**DOI:** 10.1101/2024.01.29.577879

**Authors:** Guoliang Lin, Zhiru Huang, Tingsong Yue, Jing Chai, Yan Li, Huimin Yang, Wanting Qin, Guobing Yang, Robert W. Murphy, Ya-ping Zhang, Zijie Zhang, Wei Zhou, Jing Luo

## Abstract

High-quality, chromosome-scale genomes are essential for genomic analyses. Analyses, including 3D genomics, epigenetics, and comparative genomics rely on a high-quality genome assembly, which is often accomplished with the assistance of Hi-C data. Current Hi-C-assisted assembling algorithms either generate ordering and orientation errors or fail to assemble high-quality chromosome-level scaffolds. Here, we offer the software Puzzle Hi-C, which uses Hi-C reads to accurately assign contigs or scaffolds to chromosomes. Puzzle Hi-C uses the triangle region instead of the square region to count interactions in a Hi-C heatmap. This strategy dramatically diminishes scaffolding interference caused by long-range interactions. This software also introduces a dynamic, triangle window strategy during assembly. Initially small, the window expands with interactions to produce more effective clustering. Puzzle Hi-C outperforms available scaffolding tools.

## Introduction

Analysis of genomic data rely on high-quality chromosome-level genomes. Accuracy is essential for downstream genomic analyses, and especially for 3D, comparative, and functional genomic analyses. For example, due to significant interactions being calculated on the linear distance of the genome, 3D analyses can create an incorrect assembly, which leads to false positive interactions, resulting in unreliable results(1,2). Chromosome evolution and estimation of recombination rely on the contiguity of the genome(3,4). Only chromosome-scale genomes can drive an understand the complexity of regulatory architecture and how cis-regulatory elements influence genes, because a cis-regulatory element is likely more than 1 Mb apart from the target gene(5–7). A contiguous genome can significantly improve the interpretation of genome-wide association studies (GWAS)(8–11) because usually regions of linkage are not on different contigs. Thus, high-quality chromosome-level genomes are requisite for multiple downstream genomic analyses(12).

Long-read sequencing technologies have facilitated genome assembly because they yield large, overlapping repetitive regions. Notwithstanding, such contigs do not always stretch into a complete chromosome or even one arm of a chromosome. To obtain chromosome-scale scaffolds, various strategies have been explored to increase the contiguity of *de novo* genome assemblies. Two primary methods order and orient scaffolds for chromosome-level assembly: genetic mapping and high-throughput chromatin conformation capture (Hi-C). Traditional genetic mapping orders and orients scaffolds based on linkage groups. However, the construction of a genetic map requires a large number of individual offspring to be sequenced. This dramatically limits the application of genetic maps in the genome because many species have K reproductive strategies that cannot satisfy the requirement by having enough offspring(13–15). Further, the sequencing of a large number of individuals costs time, computing resources, storage resources, and other expenses(16). By contrast, recently developed Hi-C provides a powerful tool for chromosome-level assembly. Hi-C merely requires a small number of tissue samples to mount more than 85% of sequences to chromosomes(17,18). Consequently, Hi-C is the most commonly used method for scaffolding at the chromosome level.

Three invariant features of Hi-C interactions are used for genome assembly(19): intra-chromosomal interaction enrichment, the random positioning of chromosomes in the nucleus, and the local smoothness of interactions reflected in the Hi-C heat map. In the Hi-C interaction matrix, a locus tends to interact more frequently with another locus within the same chromosome (cis-interactions) than with loci on a different chromosome (trans-interactions). Two phenomena of 3D chromatin may contribute to this feature. In the first phenomenon, chromosomes occupy distinct volumes throughout the cell cycle, leading to physical separation between chromosomes(20,21). The second phenomenon relies on the random positioning of chromosomes in the nucleus(22), which may largely reduce chromosomal interactions, and the probability of intra-chromosomal interaction decreases with increasing linear distance. Thus, in Hi-C interaction maps, the frequency of interaction tends to decrease with genomic distance, i.e., a locus interacts more frequently with other loci that are nearby in the genomic space than with distant loci. When the distance is greater than 100kb, the probability of interaction is about 1/x, where x is the distance between two points(17,23). Finally, local smoothness of interactions as reflected in the Hi-C heat map interaction of adjacent points tends to be consistent(24). Available software commonly use features one and two for scaffolding.

Accompanying the sharply decreasing price of genomic sequencing, Hi-C data for scaffolding at chromosome-level is now accessible and popular. The first Hi-C scaffolding software, LACHESIS(17), developed in 2013, has seen increasingly employment for constructing chromosome-level in genomes. Several software options exist for Hi-C scaffolding(17,25–28), however none can eliminate errors including artificial relocations, translocations, and inversions(26,28), which result in false assemblies and erroneous genomes. Chromosomes may fold into various structures, such as loops, topological associated domains (TADs), and compartments, which lead to many long-range interactions. Such interactions violate the assumption that the probability of the intra-chromosomal interaction decreases with the linear distance, thus causing an incorrect assembly. Some methods use a contig-end solution(29,30), but the employment of limited information results in disappointing performance. To address these issues, we offer an easy-to-use Hi-C scaffolding software, Puzzle Hi-C, which uses a triangle window and iterative assembly strategy to reduce long-range interactions thereby improving performance. This software achieves outstanding scaffolding results in simulated data and real data. The source code and documentation of Puzzle Hi-C are available at GitHub (https://github.com/linguoliang/puzzle-hi-c.git).

## Methods

### Datasets

We used human’s Hi-C data from the GM12878 cell line as benchmark data. We used Hi-C data from a gayal (*Bos frontalis*) and puffer fish (*Takifugu bimaculatus*) as examples. We also used the Hi-C data from a broomcorn millet (*Panicum miliaceum*), Indian cobra (*Naja naja*), Peking duck (*Anas platyrhynchos*), water buffalo (*Bubalus bubalis*), yellow croaker (*Larimichthys crocea*), and fighting fish (*Betta splendens*) (S1 Table).

### Puzzle Hi-C pipeline

The Puzzle Hi-C pipeline contains three steps: mapping, scaffolding, and building. Briefly, Puzzle Hi-C uses juicer software for mapping step. Next, scaffolding takes each chromosome group as input and iteratively merges all chromosome groups’ contigs into one chromosome. Finally, building reconstructs each chromosome by concatenating the contigs, adding gaps between the contigs, and generating the genome in the FASTA format (Fig 1).

### Puzzle Hi-C mapping

Puzzle Hi-C uses juicer software for mapping. Juicer uses bwa mem default parameters for mapping, after mapping juicer program will filter the duplicate matched sequences, in juicer it is considered that if two pairs of double-end sequence matching results, their position information only differ by 4 pb, then it is considered a duplicate sequence, juicer will remove this part duplicate sequences and keep only the result on one pairwise comparison.

### Puzzle Hi-C scaffolding

The iterative algorithm used for scaffolding solves two problems: “ordering” assigns a relative position to each scaffold on each chromosome with respect to the other scaffolds assigned to the same chromosome; and “orienting” determines which of the two ends of each scaffold is adjacent to the preceding scaffold, and which end is adjacent to the next scaffold. In each step, subsets of the input scaffolds are ordered and oriented with respect to one another to create a new, longer set of scaffolds, which are then used as inputs for the next step until all the chromosome contigs are scaffolded in one scaffold. The software uses a weight matrix to build a graph. The graph nodes are the scaffolds in a chromosome-group. Weight is defined as follows:

Each end of each scaffold is labeled using B (begin) and E (end). Given two scaffolds i and j, there are four possible connections, BB, BE, EB, and EE. We defined a length cutoff of l and considered the read pairs mapped in the region of length l at both ends (B and E) of the scaffolds.

The number of links was determined using:

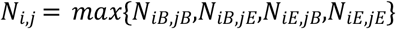

For each scaffold i, we only considered the top 5 linked edges. Next, we obtained a link-score for each pair as follows:

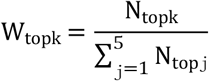

When ordering the scaffolds, each node could only have two edges, so we retained the two edges with the largest and second-largest weight. We set a cutoff for the weight; if the weight was less than the cutoff, then the connection was considered unreliable and removed. The graph found the path with all nodes>1, and constructed new scaffolds according to the path and direction of connection for the next iteration, and increased the length of l at the same time. The iteration stopped only if the number of scaffolds equaled to desired number of chromosomes.

### Puzzle Hi-C building

Once scaffolding is completed, Puzzle Hi-C builds a chromosome-level genome. Scaffolds link with 100 bp N gaps (N can be configured with Puzzle Hi-C parameters). Puzzle Hi-C also generates an agp file to record how scaffolds are assembled with the position and direction information.

### Genomic collinearity

The genomic collinearity analysis between genomes were completed using NUCMER from the MUMmer package v3.23 with default parameters(33). After alignment, we sorted the scaffolds according to collinearity to draw the final collinearity figures.

### Scaffolding error statistics

To compare the performance of each software, we aligned the genome assembly to their respective reference genomes using the program nucmer with default parameters. Alignment quality was assessed using dnadiff(33), a MUMmer utility that provides detailed information on the differences between two genomes. To get a reliable result, we sampled scaffolding 25 times, where each sample contained 5 chromosomes.

### Hi-C scaffolding in LACHESIS, SALSA2, 3D-DNA, and ALLHiC

Hi-C reads were mapped using bwa with default parameters. The SAM file, which was generate by bwa, was filtered using PreprocessSAMs.pl.

For LACHESIS, we used default parameters except CLUSTER_N, which depended on how many chromosomes should be clustered.

For SALSA2, the minimum input files were provided with the following command line:

python run_pipeline.py -a seq.fasta -l seq.fasta.fai -b alignment.bed -e GATC -o scaffolds.

For ALLHiC, we used the default parameters, except -k parameter. The k parameter was set according to how many chromosomes were clustered.

For 3D-DNA, Hi-C reads were aligned by juicer software using default parameters. Scaffolds >15 kb were retained, and the haploid model was selected in the 3D-DNA pipeline.

## Results

### Overview of the Puzzle Hi-C algorithm

Comparative analysis of existing software found that LACHESIS and ALLHiC used of all interaction information between scaffolds for clustering, and achieved high performance on clustering(17,27). However, SALSA2 only uses partial information at both ends of the scaffold, which advances ordering and orientation(28). Taking accuracy of scaffold ordering and clustering into account, Puzzle Hi-C dynamically changes the size of the statistical window at both ends of the scaffold, and increases the window size as the number of iterations increases(Fig 1). Therefore, our software uses local information at both ends of the scaffold for ordering and orientation at the initial assembly, and as the statistical window increases, it uses global interactions for better clustering (Fig 2). Puzzle Hi-C contains three steps: mapping, scaffolding, and building.

Mapping uses the Juicer software(31), which filters out duplicate, abnormal alignment, and restriction site information. scaffolding adopt an iterative method to obtain accurate assemblies via multiple iterations. Finally, the genome is assembled according to the ordering and orientation results and output in final fasta and apg format files.

**Fig 1.**
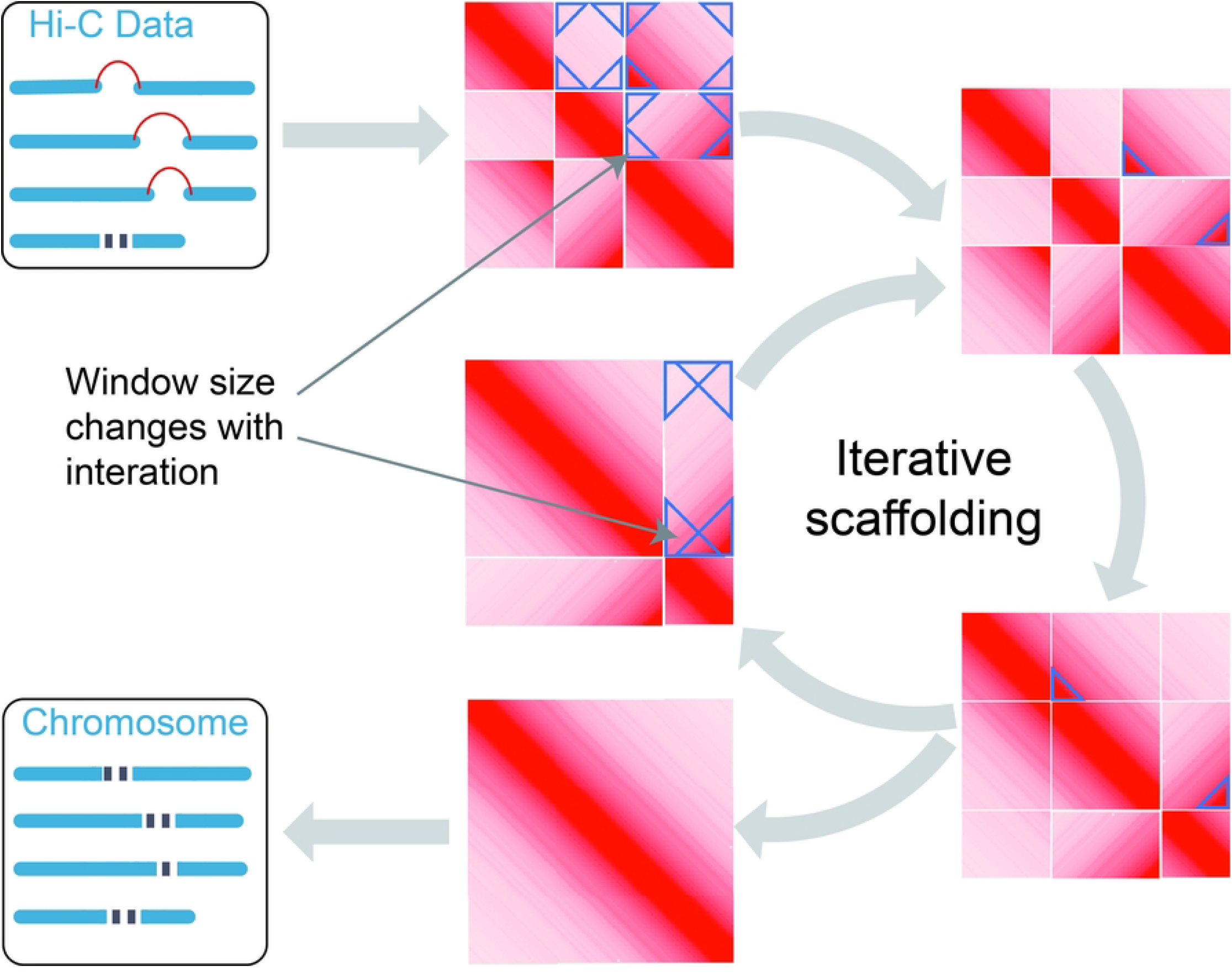
Puzzle Hi-C Pipeline. The Puzzle Hi-C pipeline contains three steps: mapping, scaffolding, and building. Ordering and orientation adopt an iterative method to obtain accurate assemblies via multiple iterations. Puzzle Hi-C introduces a dynamic, triangle window strategy during assembling. The triangle window is initially small and expands with interactions to produce more effective clustering. Finally, the genome is assembled according to the scaffolding results and output in final fasta and apg format files.

**Fig 2.**
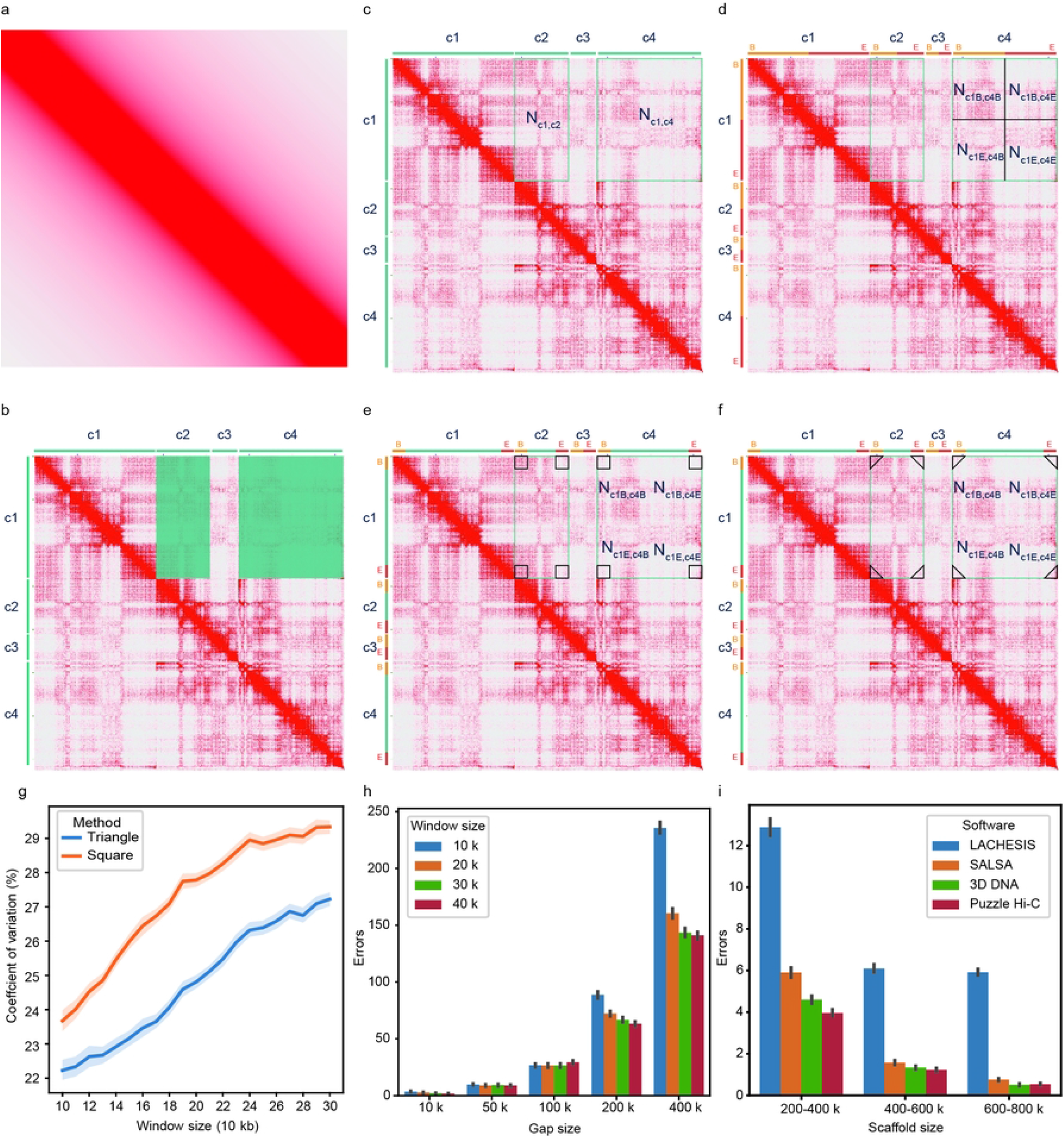
Contact probability and strategies to evaluate distance between scaffolds adopted by different software. **a**, ideally distribution of Hi-C contact, Hi-C distribution in line with 1/x. Heat map shows the diagonal position of interaction density is very high, the farther away from the diagonal, the lower the interaction density is. b, Heat map shows the chr2 [0-35MB] assembled by LACHESIS, where c1, c2, c3 and c4 represent scaffolds and the rectangle represents the number of Hi-C reads links with two scaffolds. Due to Compartment and TADs, there are many long-range interactions, which make long range interaction densities is higher than adjacent interaction densities. c, strategy to evaluate distance adopted by LACHESIS and ALLHiC. **d**, strategy to evaluate distance adopted by 3D DNA. **e**, strategy to evaluate distance adopted by SALSA. **f**, strategy to evaluate distance adopted by Puzzle Hi-C. **g**, the CV of interaction density with triangle region or square region. **h**, errors of the distance between two scaffolds with different gap size in Puzzle Hi-C. **i**, errors while different strategies to evaluate distance between two scaffolds with 1000 samplings.

### Evaluation using simulated and real data

To compare the performance of Puzzle Hi-C and other software, we used the human genome hg38. We assessed the autosomes using lengths of 200 kb, 600 kb and 1 Mb contigs.

First, we obtained statistics on the assembly results of different Hi-C scaffolding software in 200 kb, 600 kb, 1 Mb scaffolds (S2-6 Table). For example, LACHESIS assembled a genome size of 2.77 Gb and contained 102, 37 and 33 scaffolds, respectively. Scaffold N50s tended to be stable at about 135 Mb, while scaffold N90s were 79.8 Mb, 68.8 Mb, and 77.5 Mb, respectively (Supplementary Table 2). SALSA2 assembled scaffolds of 1193, 593, and 538, respectively. Scaffold N50s were 8.6 Mb,9.3 Mb, and 10.0 Mb, respectively (S3 Table). The assembled scaffolds were relatively short and the clustering effect was not ideal, with scaffold N90s of only 1.2 Mb, 2.5 Mb, and 2.8 Mb, respectively. The genome assembled by Puzzle Hi-C was quite similar to that of LACHESIS. The assembled scaffolds were 703, 191, and 109, respectively, and scaffold N50s were 128.5 Mb, 130.9 Mb, and 154.4 Mb, respectively (S6 Table). Scaffold N90s were 44.8 Mb, 55.6 Mb, and 56.6 Mb, respectively. Genome assembly size, scaffold N50, and scaffold N90 can only reflect the clustering effect of Hi-C-based scaffolding software. Because LACHESIS clusters before assembly, the Scaffold N50 result was excellent.

Second, to evaluate and compare the performance of existing scaffolding software in scaffolding and orientation, we used dnadiff to compare the quality of the genomes. We assessed three features: the number of relocations as determined by the number of breaks in the alignment of scaffolds belonging to the same chromosome, but not consistently ordered; the number of translocations, that being the number of breaks in the alignment of scaffolds belonging to different chromosomes; and the number of inversions, or breaks in the alignment by scaffolds inverted with respect to one another. As the size of scaffolds got smaller, the proportion of assembly errors in LACHESIS increased (Table 1). It produced 69 assembly errors in the 1 Mb scaffold size, including 9 translocations, 31 orientation assembly errors, and 29 relocations; at the 600kb scaffold size, assembly errors increased to 132, and the 200kb size had 380 errors, which showed an inverse relationship between scaffold size and assembly errors. Other software showed the same pattern. Comparatively, Puzzle Hi-C consistently achieved the greatest assembly accuracy under different sizes of scaffolds (Fig 3a-c, Table 1), for example having 67 assembly errors at the 1 Mb scaffold size (relocations 11, translocations 8, inversions 48).

**Table 1.**
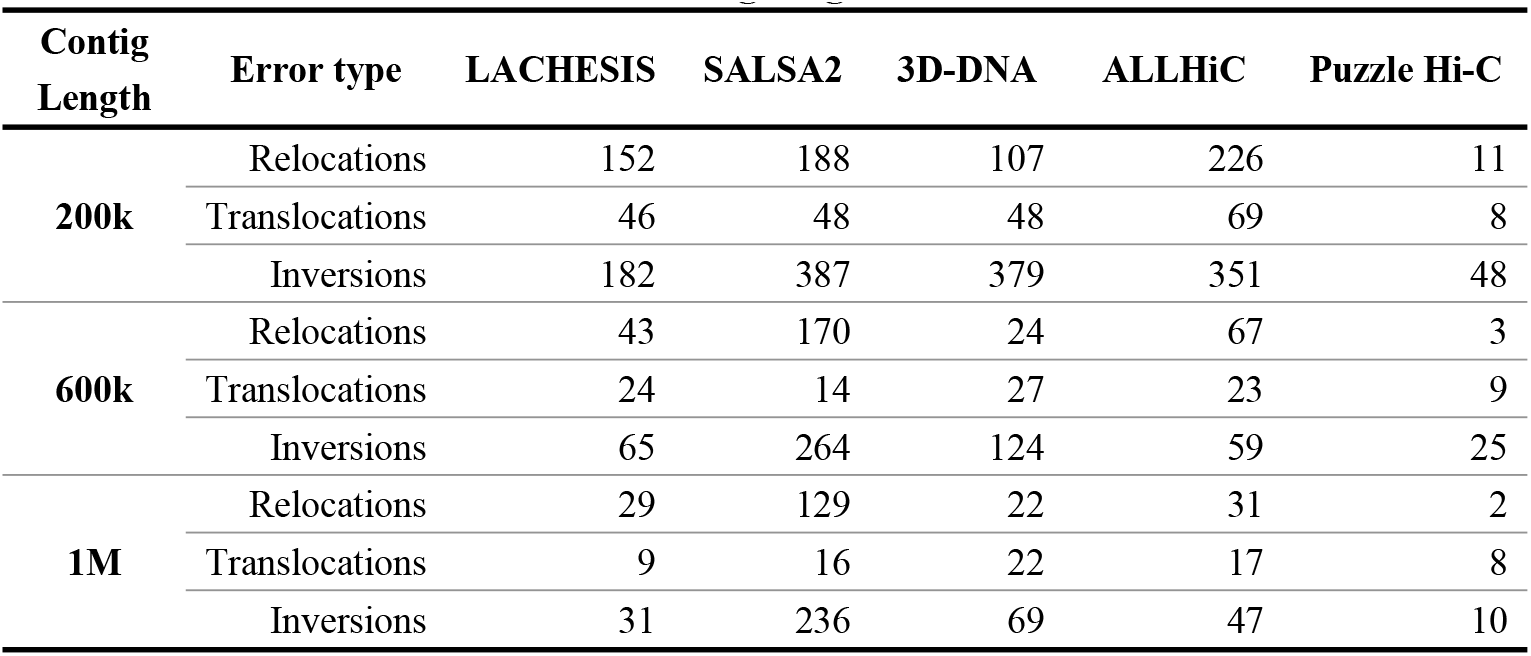
Statistical errors generated by different software with different contig length.

| Contig Length | Error type | LACHESIS | SALSA2 | 3D-DNA | ALLHiC | Puzzle Hi-C |
| --- | --- | --- | --- | --- | --- | --- |
| 200k | Relocations | 152 | 188 | 107 | 226 | 11 |
|  | Translocations | 46 | 48 | 48 | 69 | 8 |
|  | Inversions | 182 | 387 | 379 | 351 | 48 |
| 600k | Relocations | 43 | 170 | 24 | 67 | 3 |
|  | Translocations | 24 | 14 | 27 | 23 | 9 |
|  | Inversions | 65 | 264 | 124 | 59 | 25 |
| 1M | Relocations | 29 | 129 | 22 | 31 | 2 |
|  | Translocations | 9 | 16 | 22 | 17 | 8 |
|  | Inversions | 31 | 236 | 69 | 47 | 10 |

**Fig 3.**
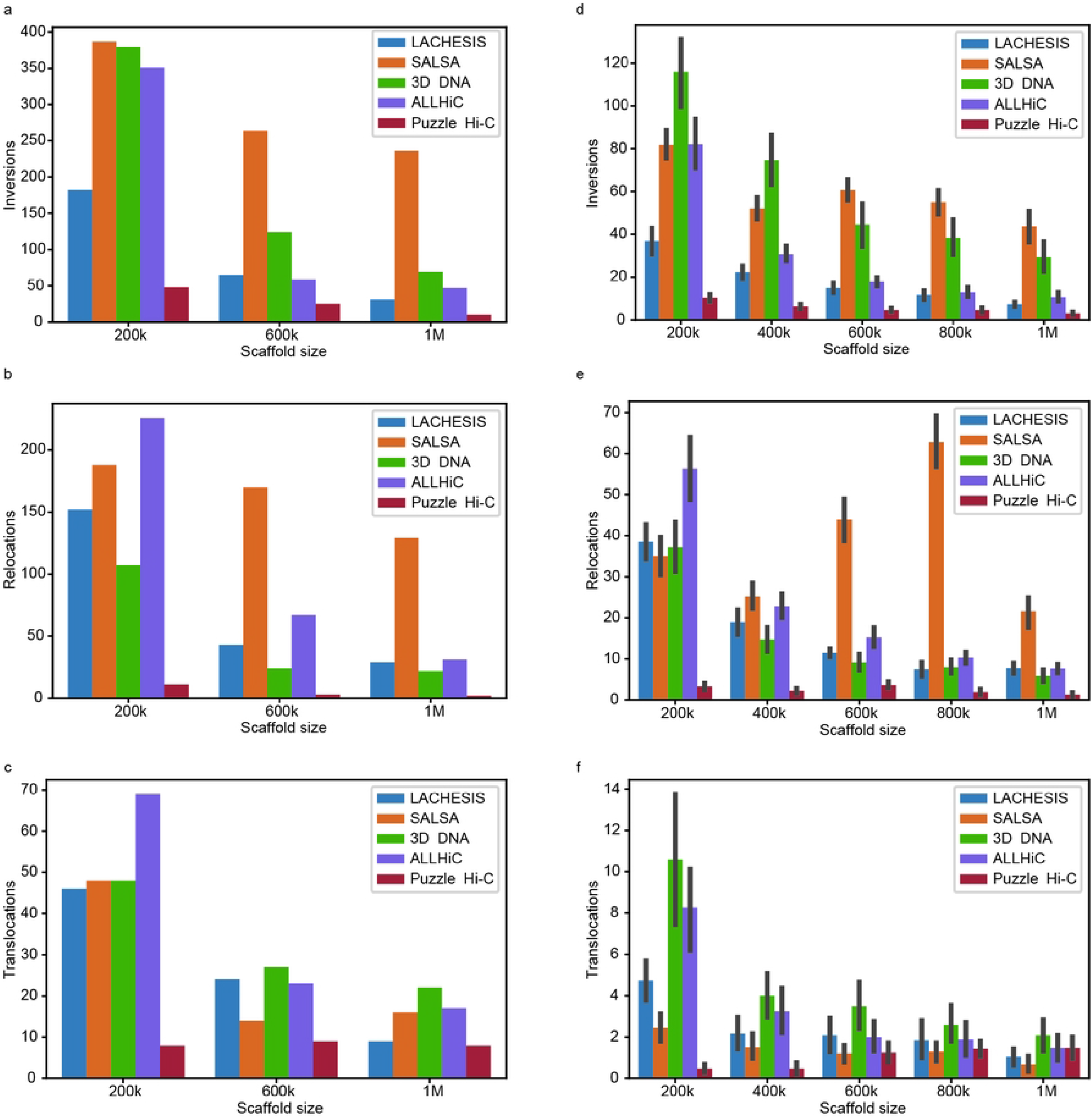
Statics different errors generated by different software with different scaffold size. **a-c**, inversions, relocations and translocations generated by LACHESIS, SALSA, 3D DNA, ALLHiC and Puzzle Hi-C under different length of Scaffolds. **d-f**, inversions, relocations and translocations generated by LACHESIS, SALSA, 3D DNA, ALLHiC and Puzzle Hi-C with 25 sampling data under different length of Scaffolds.

Third, we resampled the human genome 25 times with five different scaffold sizes. Puzzle Hi-C outperformed the other software packages (Fig 3d-f). For example, it produced the best chromatin assembly, showing much less assembly errors. Puzzle Hi-C was not affected by the size of the scaffolds, and it was more robust (Fig 3d-f).

To test the assembly performance of Puzzle Hi-C using real data, we also employed the scaffold version of the human genome assembly (version:GCA_001013985.1). This analysis used LACHESIS. We compared the assemblies to the human genome GRCh38 using MuMmer software. LACHESIS produced 999 errors in its ordering and orientation of large fragments in assembly, and Puzzle Hi-C gave 647 errors, except for chromosome 1, which was composed of three scaffolds. In addition to assembly errors of large fragments, LACHESIS also had more problems in assembling small scaffolds, such as chromosomes 17, 19, 20, and 22. Puzzle Hi-C did not exhibit this problem (Fig 4, Table 2). Other methods also showed more errors than Puzzle Hi-C (Table 2). We also assembled other species genomes across a range of taxonomic groups, genome sizes and initial assembly quality. Puzzle Hi-C consistently generated assemblies with higher contiguity (Supplementary Figure 1).

**Fig 4.**
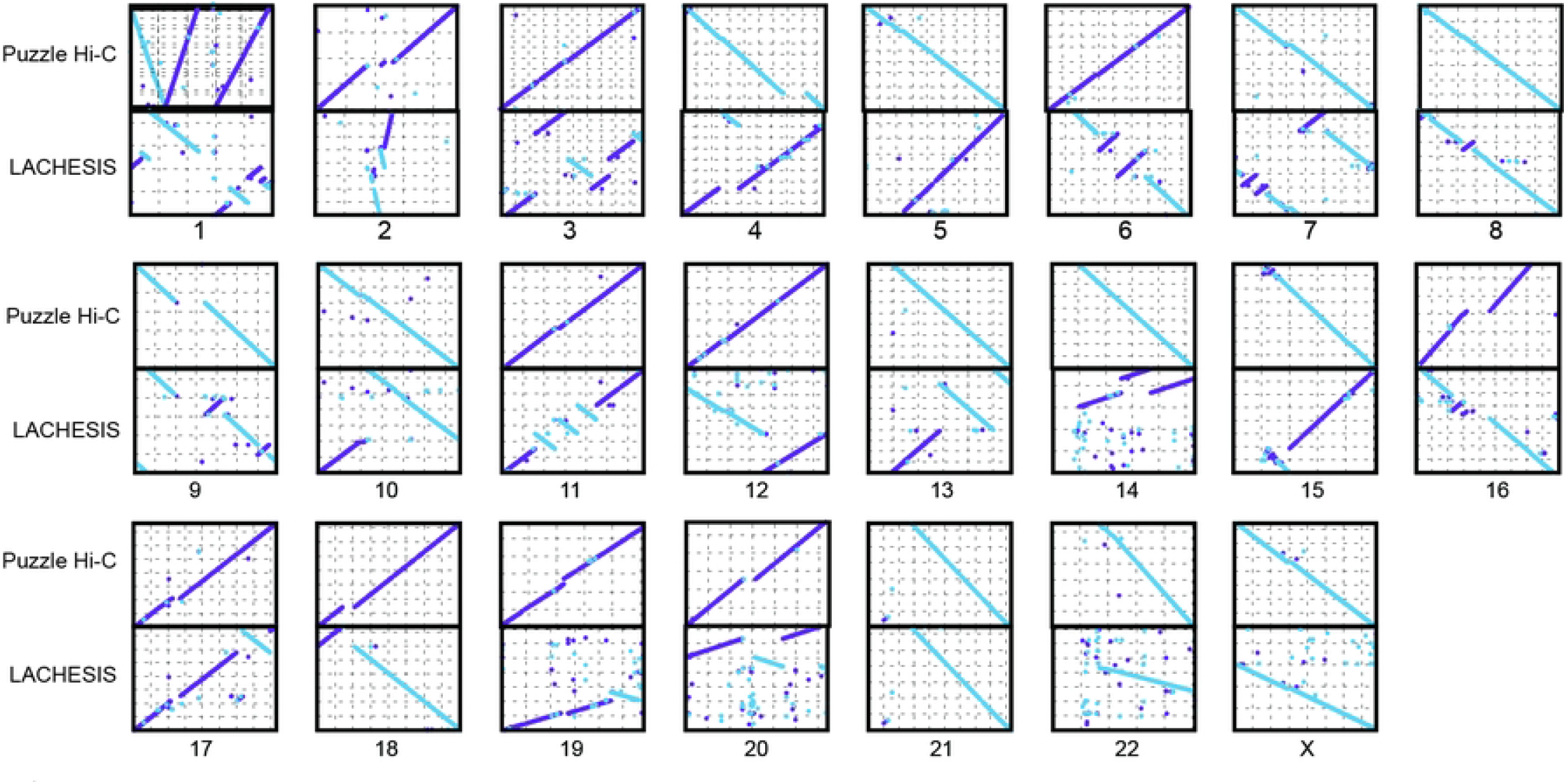
The synteny of chromosomes assembled by Puzzle Hi-C and LACHESIS compared with GRCh38.

**Table 2.**
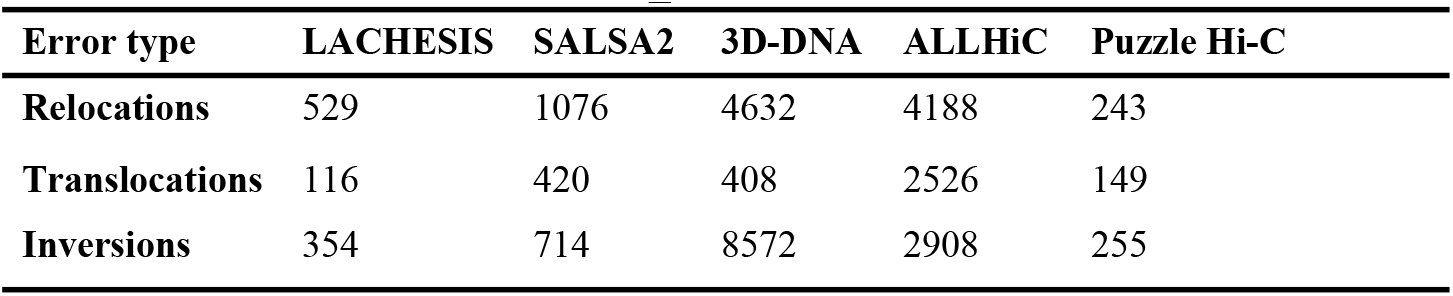
Statistical errors generated by different software with GCA_001013985.1.

### Assembling genomes enriched with long-range interactions

To further test the robustness of assembly by Puzzle Hi-C, we employed chromosome 2 of gayal, which contains a Robertsonian translocation. This chromosome has more repetitive sequences and long-range interactions than its relatives. Long-range interaction will obstruct ordering prediction. The Hi-C interaction matrix revealed a very strong internal interaction of compartments on chromosome 2, which indicated a long-distance interaction. Other software assemblies also detected the rearrangement of large fragments, but Puzzle Hi-C scaffolding software obtained relatively fewer chromatin orientation assembly errors. Therefore, Puzzle Hi-C appeared to best assemble chromosomes when chromatin interactions occurred (Fig 5).

**Fig 5.**
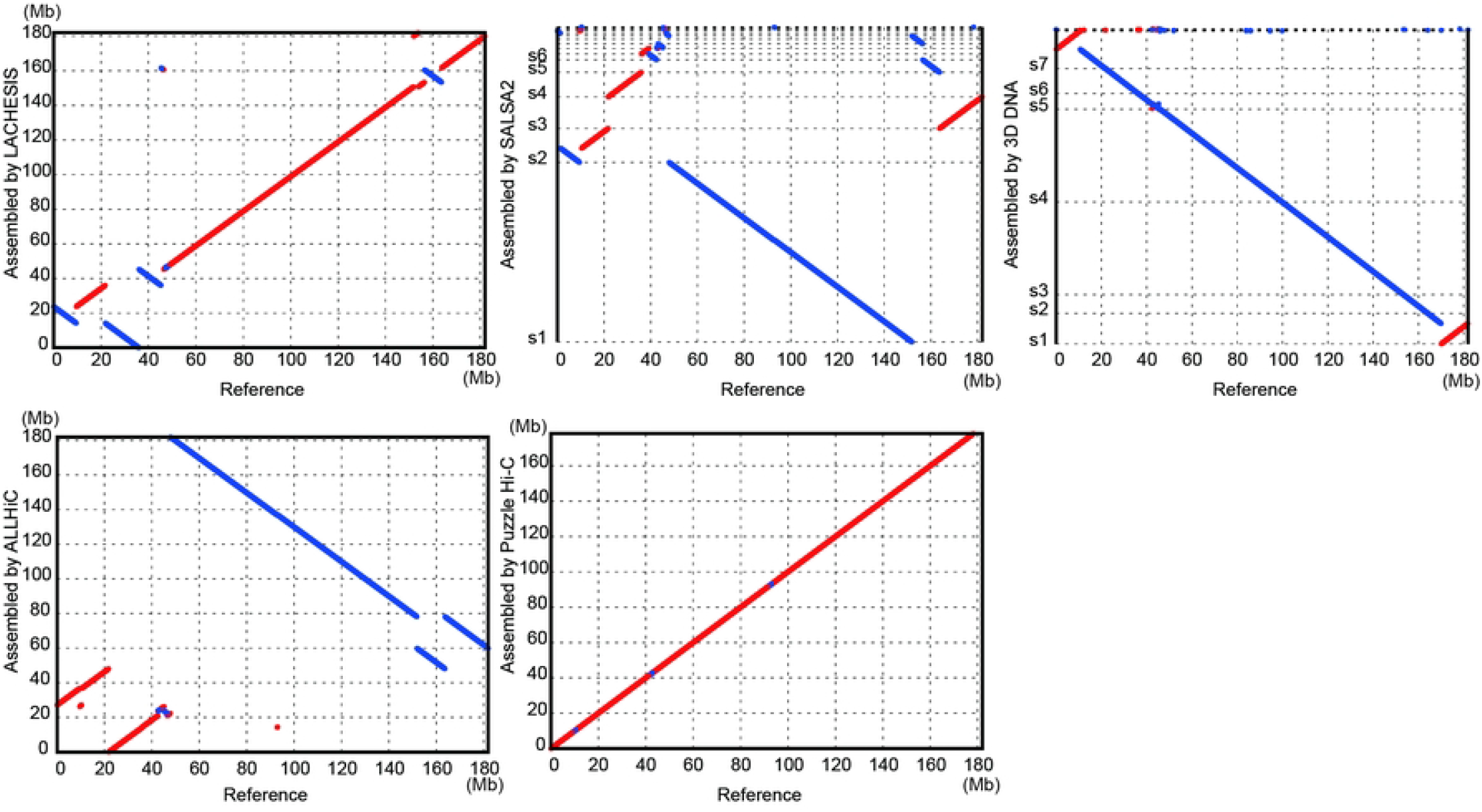
The synteny of chromosomes assembled by Puzzle Hi-C and other software compared with gayal chromosome 2.

### Genome’s quality is curial for 3D analysis

A high-quality genome is essential for downstream analysis. However, due to the absence of reliable tools, chromosome-level assemblies may contain some error. These errors may directly affect the main results of the analysis. To estimate the effect of chromosome errors on downstream analysis, we downloaded the genome of *Takifugu bimaculatus*(32), which was assembled by LACHESIS. Compared to the genome of *T. rubripes*, the genome of *T. bimaculatus* has 809 inversions and 2618 relocations. We reassembled this genome using Puzzle Hi-C and obtained 519 inversions and 1791 relocations. We performed comparable analysis on both old and new genomes. The results showed that different genome assembles will affect comparative analysis (Fig 6).

**Fig 6.**
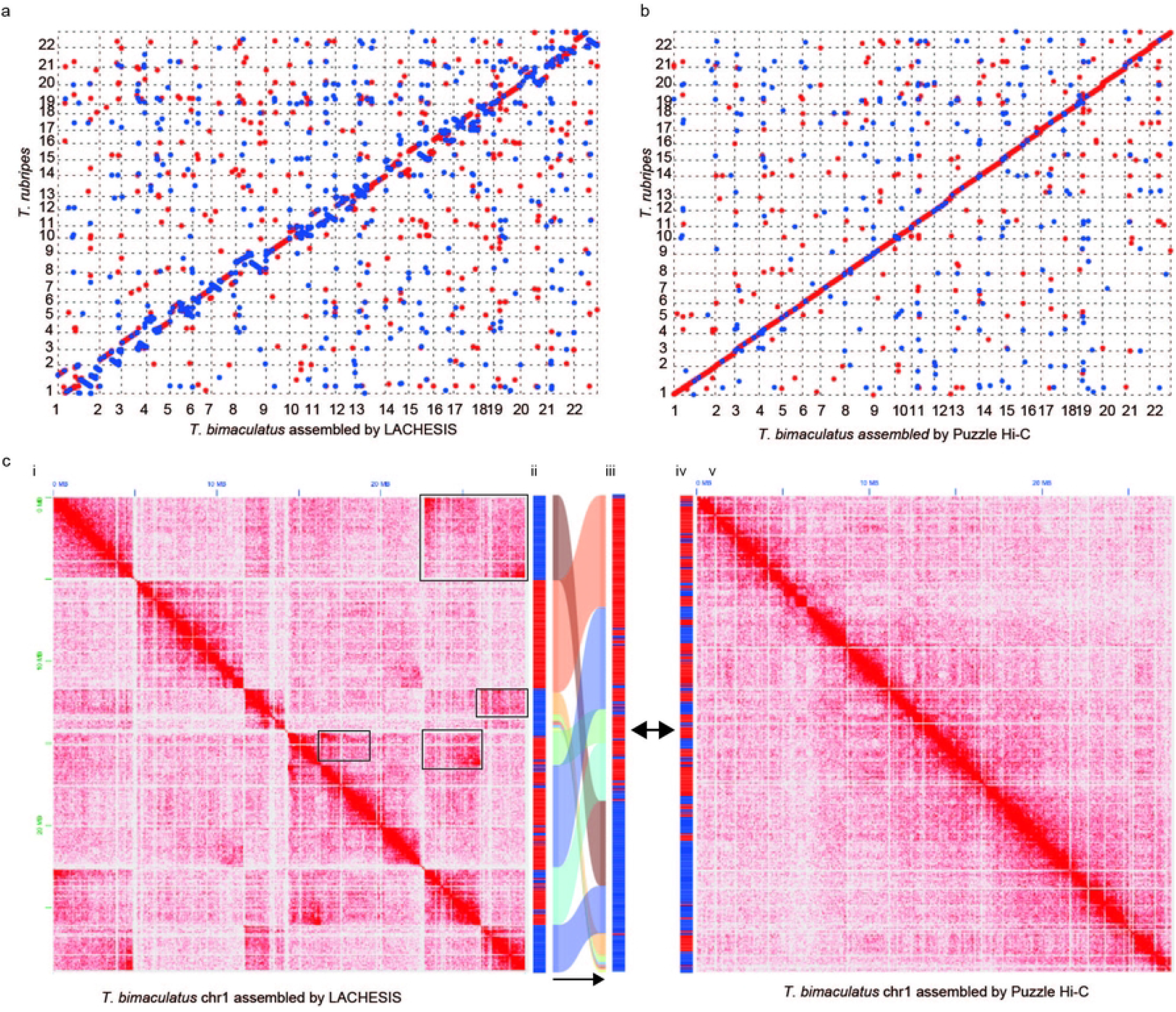
The scaffolding result of LACHESIS and Puzzle Hi-C on *T. bimaculatus*. **a**, the syteny between *T. bimaculatus* and *T. rubripes*; **b**, the syteny between Puzzle Hi-C corrected *T. bimaculatus* and *T. rubripes*; **c**, i *T. bimaculatus* genome chr1 Hi-C heat map, the black box is the Hi-C heat map suggesting assembly error; ii *T. bimaculatus* genome chr1 Compartment, red is Compartment A and blue is Compartment B; iii is the rearrangement of *T. bimaculatus* genome chr1 Compartment according to the corrected chr1; iv Puzzle Hi-C corrected chr1 Compartment after Puzzle Hi-C correction; v Puzzle Hi-C corrected chr1 Hi-C heat map.

## Discussion and Conclusion

Hi-C data facilitate the assembly of chromosome-level genomes by locating and avoiding long-range interactions. Puzzle Hi-C uses a dynamics triangle window to calculate interaction densities. It dynamically changes the size of the triangle at both ends of the scaffold. Windows start small in initial iterations, which facilitates the assembling of smaller scaffolds. Puzzle Hi-C excludes long-range interactions when the window is small. While LACHHESIS(17), 3D DNA(25), and ALLHiC(27) use all interaction information between two scaffolds, such can result in errors in ordering due to long-range interactions. As iterations increase in Puzzle Hi-C, the window increases in size, therefore obtaining better chromosome clustering by selectively using all interaction information. Such avoids the problem of failing to cluster scaffolds into chromosomes when using only local interaction information, which SALSA(26,28) does. Puzzle Hi-C evaluation on human genome outperforms ALLHiC(27), LACHESIS(17), 3D DNA(25), and SALSA(26,28) in ordering and orientation in both simulated and real data, and with robust performance. The same result occurs upon applying Puzzle Hi-C to all other tested genomes. Further, Puzzle Hi-C outperforms other software when assembling the complex gayal genome, which has many long-range interactions. Similarly, the reassembled genome of the puffer fish reveals improvements when compared with the original assembly(32). Finally, the results suggest that the genome-quality greatly impacts 3D genome analysis. Thus, accurate 3D genome analysis requires accurate chromosome-level genomes.

## Data and Code Availability

All Hi-C data were downloaded from NCBI (S1Table). The Puzzle Hi-C software package with a detailed user tutorial and sample input and output files can be found at https://github.com/linguoliang/puzzle-hi-c.git.

## Acknowledgments

G.L., Z.Z., Y.Z., W.Z., R.W.M., and J.L. conceived and designed the study. G.L. developed the method with help from Z.H., T.Y., J.C., Y.L., Z.H., T.Y., H.Y., W.Q., and G.Y. performed analysis. G.L., Z.H. and T.Y. wrote the manuscript. G.L., Z.H., T.Y., Z.Z., R.W.M., W.Z., and J.L. revised the manuscript. Computational resources were provided by the Advanced Computing Center of Yunnan University.

## Supporting information

S1 Text. Puzzle Hi-C pipeline.

S1 Fig. Hi-C contact maps of assemblies constructed from puzzle hic.

S1 Table. SRA data used in this study.

S2 Table. LACHESIS scaffolding results.

S3 Table. SALSA2 scaffolding results.

S4 Table. 3D DNA scaffolding results.

S5 Table. ALLHiC scaffolding results.

S6 Table. Puzzle Hi-C scaffolding results.

## Notes

### Competing Interest Statement

The authors have declared no competing interest.

